# Hybrid-derived weedy rice maintains adaptive combinations of alleles associated with seed dormancy

**DOI:** 10.1101/2022.03.15.484373

**Authors:** Toshiyuki Imaizumi, Yoshihiro Kawahara, Gabriela Auge

**Affiliations:** Institute for Plant Protection, National Agriculture and Food Research Organization (NARO), Tsukuba, 305-8604, Japan; Research Center for Advanced Analysis, NARO, Tsukuba, 305-8518, Japan; Consejo Nacional de Investigaciones Científicas y Tecnológicas (CONICET) - Instituto de Biociencias, Biotecnología y Biología Traslacional, Facultad de Ciencias Exactas y Naturales, Universidad de Buenos Aires, Buenos Aires, Argentina

**Keywords:** hybridization, genome stabilization, adaptive introgression, seed dormancy, weedy rice

## Abstract

Hybridization is a widespread phenomenon in plants and is a pathway for the evolution of adaptive traits. However, this process may also affect the persistence of combinations of adaptive alleles evolved through natural selection when hybridization occurs between adapted and non-adapted populations. Hybridization between weedy and cultivated rice has been confirmed with an adaptive introgression of deep seed dormancy alleles from cultivated rice. In this study, we explored the influence of hybridization on the conservation of combinations of adaptive alleles by evaluating the natural variation in and the genetic structure of genomic regions associated with seed dormancy. Based on sequence variation in the genomic regions associated with seed dormancy, we revealed that hybrid-derived weedy rice strains maintained most of the adaptive combinations for this trait that were observed in the parental weedy rice, despite equal representation of the parental weedy and cultivated rice in the whole genome sequence. Moreover, the hybrid-derived weedy rice strains had deeper seed dormancy than their parental weedy rice strains. This study suggests that hybridization between weedy rice (having adaptive allelic combinations for seed dormancy) and cultivated rice (having non-adaptive combinations) generates weedy rice strains that express deep seed dormancy caused by genome stabilization through the removal of alleles derived from cultivated rice, in addition to the adaptive introgression of deep seed dormancy alleles derived from cultivated rice. Thus, hybridization between adapted and non-adapted populations seems to be reinforcing the trajectory towards the evolution of adaptive traits.

## 1 INTRODUCTION

The factors that maintain genetic and phenotypic variation within natural populations have long been of interest to evolutionary biologists as they provide clues on the factors influencing local adaptation (Delph & Kelly, 2014). Identifying the genetic mechanisms underlying adaptation remains an important pursuit in evolutionary biology. Strong evidence demonstrates that genetic and phenotypic variation facilitates establishment success and population persistence in novel and/or unpredictable environments (Forsman, 2014). One important source for genetic variation is hybridization, which is a widespread phenomenon in plants (Arnold, 1992; Ellstrand et al., 1996; Rieseberg et al., 2007). Hybridization with non-adapted populations (or genetically distinct populations) may break their effective gene combinations; however, these combinations might also provide a selective advantage even after having been partially broken up after hybridization due to transgressive traits in hybridizing populations (Anderson, 1953; Rieseberg et al., 1999; Ellstrand, 2003). There is limited knowledge of how the effective combinations of adaptive alleles have evolved through natural selection and have been maintained despite hybridization with non-adapted populations (Todesco et al., 2020).

Weedy rice (*Oryza* spp.) is an especially good study model for revealing the effects of hybridization on the effective combinations of adaptive alleles given its very close relationship to the genomic model cultivated rice (*O. sativa* L.) and its set of wild-like traits. In general, weedy rice has evolved independently multiple times from domesticated ancestors, and this evolution has been confirmed from four different cultivated rice subgroups: *temperate japonica* (hereafter, TEJ), *tropical japonica* (TRJ), *indica* (IND) and *aus* (AUS) (Li et al., 2017; Qiu et al., 2017; Vigueira et al., 2019; Sun et al., 2019; Qiu et al., 2020; Imaizumi et al., 2021). In Japan, two major morphotypes exist: black-hull (BH) and strawhull (SH); these morphotypes are classified based on hull colour. Whole genome sequence analyses have revealed that Japanese weedy rice can be classified into three groups: TEJ-derived BH weedy rice, TEJ-derived SH weedy rice, and TRJ-derived SH weedy rice (hereafter BH, SH_TEJ and SH_TRJ, respectively), with SH_TEJ weedy rice evolving from the hybridization between BH weedy rice and cultivated rice (Imaizumi et al., 2021). The hybrid-derived weedy rice strains display crop-like adaptive traits, such as shorter plant height, which facilitates crop mimicry, as well as a critical weedy-like trait, namely deep seed dormancy. Cultivated rice is expected to be a non-adaptive population for weedy-like traits, including seed dormancy, so the system provides a unique opportunity for evaluating how the effective combinations of adaptive alleles are maintained after hybridization with non-adapted populations.

Seed dormancy is a major life-history trait crucial for local adaptation because it regulates the timing of germination and determines the post-germination environment (Donohue et al., 2010). However, the factors that maintain variation in seed dormancy remain largely unknown. The dormancy level of seeds at the time of dispersal (primary dormancy) is established during maturation in maternal plants, and it regulates the timing of germination. Induction of primary seed dormancy is regulated by genes, the maternal environment and their interaction, and the process is influenced by many quantitative trait loci (QTLs) and a wide range of environmental factors. In rice, at least 30 QTLs have been found for seed dormancy, but only three of them have been subjected to map-based cloning to identify the underlying genes: *Seed dormancy 4* (*Sdr4*, Sugimoto et al., 2010); *SD7-1/Rc* (Gu et al., 2011); and *Seed dormancy1-2* (*qSD1*, Ye et al., 2015), also known as *semidwarf-1* (*sd-1*). In addition, *OsG*, the orthologue of soybean *G*, has been shown to be responsible for seed dormancy based on analysis of transgenic plants carrying an overexpression construct with this gene (Wang et al., 2018). Although seed dormancy is regulated by a large number of genes, it is an extremely plastic trait that is influenced by a wide range of environmental cues. Plasticity in dormancy is modulated both by pre-and post-dispersal environmental cues such as temperature, light and, to a lesser extent, nitrate. Temperature during seed maturation can be considered the predominant environmental signal that controls post-dispersal primary dormancy levels. Specifically, low temperatures during seed maturation dramatically increase dormancy in *Avena fatua* (Peters, 1982), wheat (Reddy et al., 1985) and *Arabidopsis thaliana* (Chen et al., 2014; Burghardt et al., 2016), whereas high temperatures increase seed dormancy in cultivated rice (Suriyasak et al., 2020). Studying the interaction of the genetic and environmental factors controlling seed dormancy may provide insights into the evolutionary trajectory of this trait in lineages of interest that affect adaptation to novel environments, such as the highly invasive and pervasive weedy rice.

Here, we evaluated the natural variation related to seed dormancy in weedy rice strains collected throughout Japan to reveal how hybridization with cultivated rice has affected adaptive combinations of alleles associated with seed dormancy. To gain insights into genomic variation, we studied sequence variation among strains in the genomic regions associated with seed dormancy. Consistent with the observed phenotypes, our genomic analyses revealed that the hybrid-derived weedy rice strains maintain adaptive combinations of seed dormancy alleles derived from the parental weedy rice strain despite their admixed ancestry between weedy rice and cultivated rice based on the whole genome sequences. Altogether our study sheds light on the effects of hybridization on the expression and maintenance of adaptive traits and the genetic basis in weedy rice.

## 2 MATERIALS AND METHODS

### 2.1 Plant materials and whole genome sequence data

We used multiple weedy rice strains collected throughout Japan and one cultivated rice strain (Koshihikari) to evaluate the natural variation in seed dormancy. The experiments were repeated four times from 2017 to 2020 using seeds collected from weedy and cultivated rice strains grown in a paddy field or containers. We selected 18 weedy rice strains (10 BH, 5 SH_TEJ, and 3 SH_TRJ) in 2017, 19 weedy rice strains (11 BH, 5 SH_TEJ, and 3 SH_TRJ) in 2018 and 2019 and 11 weedy rice strains (4 BH, 4 SH_TEJ, and 3 SH_TRJ) in 2020 to collect seeds for germination assays as described in Table S1. Seeds of weedy rice strains were sown in seedling trays with commercial rice-cultivation soil, and then, the trays were placed in a greenhouse until transplanting. Seedlings were transplanted into a paddy field, except in 2017. We used 50 cm x 50 cm x 30 cm containers with paddy field soil to grow plants in 2017. The experiments were conducted at the Tsukuba-Kannondai test field at National Agriculture and

Food Research Organization (NARO) (36.0°N, 140.1°E). A total of 356 kg per ha of chemical fertilizer (N:P:K = 14:14:14%) was basally applied before transplanting, and the plants were transplanted at one plant per hill with 15 cm x 30 cm spacing. The plants were harvested approximately six weeks after flowering, when the seeds achieved physiological maturity. After harvest, the seeds were air-dried and kept in dry storage at 20°C in a desiccator cabinet until use in the germination assays. Weedy rice strains, the schedule of rice cultivation, and the timing of harvests are described in Table S1.

To evaluate genetic variation, we identified sequence variation in the genomic regions related to seed dormancy to compare the genomic basis of seed dormancy among strains. We selected and reanalysed 144 published whole genome sequence data and variation data that were used in our previous study (Imaizumi et al. 2021). The selected sequences represent the genetic diversity of Japanese, Chinese and US weeds and Japanese crops. In summary, we selected 49 Japanese weedy rice strains (25 BH, 15 SH_TEJ, and 9 SH_TRJ strains); 46 Japanese landrace strains from the TEJ, TRJ or IND group (landrace_TEJ, landrace_TRJ and landrace_IND, respectively); and 10 Japanese modern strains from the TEJ group (modern_TEJ). The non-Japanese weedy rice strains were from the Chinese TEJ or IND group and US IND or AUS group (China_TEJ, China_IND, US_IND and US_AUS, respectively), as described in Table S2.

### 2.2 Germination assays

Seeds from 9 and 12 biological replicates (different maternal plants) of each strain were used for germination assays conducted in 2017 and 2019, respectively, and seeds from 12 technical replicates (bulked seeds collected from more than 20 maternal plants) were used in 2018 and 2020 (Table S1). We used two incubation conditions, namely, darkness at 15°C or 30°C, and temperature manipulation was used to facilitate the detection of genetic variation for seed dormancy by exposing seeds to more (30°C) and less (15°C) permissive germination conditions, thereby minimizing the chance of extreme germination proportions of 0% and 100% across all strains. Each replicate comprised 20 seeds sown in 55 mm Petri dishes with 5 ml of distilled water. The plate positions were randomized within each incubation treatment after each germination census to minimize position effects.

Germination was scored at 4, 7, 10 and 14 days after sowing, with germination indicated by the protrusion of the radicle from the seed coat. Germinated seeds were removed from the Petri dishes at each census. Germination usually reached a clear plateau by 14 days of incubation. The final proportion of viable germinated seeds was used for analysis, with each plate representing the unit of analysis. Seed viability was assessed by pressing the seeds with a pestle: viable seeds were firm and not easily broken, while inviable seeds were soft and easily crushed. If decay was observed during the incubation, that information was recorded, and seeds were removed from the dishes. To assess whether genetic variation in dormancy persisted over the course of after-ripening, we used seeds after-ripened for 1, 5 and 10 weeks for the germination assays. For seeds collected in 2019 and 2020, the assays were also conducted using 20-week-old after-ripened seeds given the limited number of germinated seeds at 15°C in the dark, even at 10 weeks after harvest.

### 2.3 Analysis of population structure for genomic regions conferring seed dormancy

We used genomic data from weedy and cultivated rice strains (Table S2) to estimate ancestry proportions of genomic regions associated with seed dormancy for individuals using NGSadmix (Skotte et al., 2013). The analysis implements a clustering method similar to that in the population program ADMIXTURE (Alexander et al., 2009). The genomic data were obtained and analysed as described by Imaizumi et al. (2021). The raw paired-end reads were first trimmed with Trimmomatic version 0.38 (Bolger et al., 2014). Then, the trimmed reads were aligned to Os-Nipponbare-Reference-IRGSP-1.0 (Kawahara et al., 2013) pseudomolecules using BWA-MEM (release 0.7.17, Li & Durbin, 2010). BAM files after removal of duplications by Picard tools (release 2.18.17, Picard_toolkit, 2019) were used for NGSadmix. The genotype likelihoods used for NGSadmix were estimated by ANGSD (Korneliussen et al., 2014) with the BAM files. We extracted genomic regions associated with seed dormancy (Table S3) from the BAM files using SAMTOOLS (Danecek et al., 2021). The genomic regions were determined using gene information with the keyword “dormancy (GO:0010162 - seed dormancy or TO:0000253 - seed dormancy)” or “germination (GO:0009845 - seed germination or TO:0000430 -germination rate)” in Oryzabase (Yamazaki et al., 2010), and genes with genomic positions were used for analysis. The whole genome sequences were also analysed by NGSadmix to compare the population structure of the whole genome with that of the genomic regions associated with seed dormancy. For NGSadmix, genotype likelihoods generated from 23.3x average genome coverage sequences and a combined genotype data set of 4,922 and 9,664,095 sites for seed dormancy and the whole genome, respectively, across 144 samples were used. NGSadmix runs were performed for *K* values ranging from 2 to 10, and the optimal *K* was estimated using the likelihood of *K* (*L(K)*, Pritchard et al., 2016) and the Δ*K* statistic (Evanno et al., 2005) as the selection criterion.

Principal component analysis (PCA) was also conducted using the genomic regions associated with seed dormancy or the whole genome to assess the genetic relationships within and among 111 TEJ-derived weeds and TEJ crop rice strains using PCAngsd ver 0.97 (Meisner & Albrechtsen, 2018). PCAngsd used the genotype likelihoods with the BAM files as with NGSadmix, as described above. A combined genotype data set of 1,921 and 3,719,401 sites for seed dormancy and the whole genome, respectively, across the 111 TEJ samples was used for all analyses with ANGSD, NGSadmix, and PCAngsd; these analyses were conducted with default parameters as described in the manuals.

### 2.4 Greenhouse experiments to confirm maternal temperature effects on progeny dormancy

To examine the effects of temperature after heading on progeny dormancy, we repeated the experiment twice in 2016 and 2018 using BH and SH_TEJ weeds. JP_AC90 from BH and JP_AC112 from SH_TEJ (described in Table S2) were used in 2016, and JP_1180 from BH and JP_1181 from SH_TEJ were used in 2018. Weedy rice seeds were sown in seedling trays with rice cultivation soil and placed in a greenhouse until transplanting. Seedlings were transplanted individually into 1/5000a Wagner pots (height 20 cm; diameter 16 cm) filled with paddy field soil, and basal application of chemical fertilizer (N:P:K = 14:14:14%) was performed at a rate of 712 mg per pot before planting. Plants were transplanted at one plant per pot. This study was conducted at the Tsukuba-Kannondai test field at NARO (36.0°N, 140.1°E), and plants were grown under natural conditions until heading. Then, they were moved into a greenhouse environment just before heading when flag leaf emergence was confirmed (one to three days before heading). The greenhouse conditions were 28°C/22°C during the day/night for the control treatment and 34°C/28°C during the day/night for the high-temperature treatment. Plants from all treatments were harvested approximately six weeks after heading, when seeds had achieved physiological maturity. After harvesting, seeds were dried and kept in dry storage at 20°C until use in the germination assays. Seeds from four or eight biological replicates (different maternal plants) of each strain were used for germination assays in 2016 and 2018, respectively. Germination was scored as described above.

### 2.5 Statistical analysis

All analyses were performed in R ver. 3.63 (R Core Team, 2020). To evaluate natural variation for seed dormancy, final germination rates (number of germinants/number of viable seeds at the last census date) were analysed with binomial generalized linear mixed models using “glmer” in the “lme4” (v1.1.26) package (Bates et al., 2015). We first used a full model that included germination rates as the dependent variable and dry storage after harvest (“After-ripening”), types of weedy rice (“Type”: BH, SH_TEJ and SH_TRJ), temperature of germination assay (“Temp”), and experimental year (“Year”) as fixed factors. We classified “After-ripening” as a continuous variable and the other factors as categorical variables. Weedy rice strains (“Strain”) were used as a random effect for “Type”. Analysis of deviance based on Chi-squared values was performed using the function “Anova” in the “car” (v3.0.10) package. We also tested for the effects of “After-ripening”, “Type”, “Year” and their interactions for each incubation temperature separately with the generalized linear mixed model as described above. To interpret differences in seed dormancy among types of weedy rice, we next tested for the effects of “After-ripening”, “Type”, “Temp”, and their interactions for each year separately with the generalized linear mixed model as described above. We visualized germination after different periods of dry storage and estimated time to 50% germination using the estimated values and 95% confidence intervals (CIs) from the model.

To evaluate the effects of temperature after heading on progeny seed dormancy using the results across a 4-year timeline for the natural variation of this trait, the final germination rates of 5-week-old after-ripened seeds under 30°C and dark conditions and those of 10-week-old seeds under 15°C and dark conditions were analysed with binomial generalized linear mixed models as described above. We tested for the effects of cumulative daily mean temperature for 40 days after heading (“Maternal”) for each type of weedy rice and the germination conditions separately using “Maternal”“ as a continuous variable and “Strain” as a random effect. We visualized the effects of the cumulative daily mean temperature on germination and estimated the odds ratios of the maternal temperature effect increased by 20°C using the estimated values and the 95% CIs from the model. Regarding the greenhouse experiments, the final germination rates were analyzed with binomial generalized linear mixed models as described above. We used a full model that included germination rates as the dependent variable and “After-ripening”, “Type” (BH and SH_TEJ), “Temp”, and maternal temperature (“Maternal-Temp”: 28°C/22°and 34°C/28°) as fixed factors. We classified “After-ripening” as a continuous variable and the other factors as categorical variables for each experimental year separately. Biological replications were used as a random effect.

## 3 RESULTS

### 3.1 Natural variation in seed dormancy in weedy rice from Japan

To assess natural variation in seed dormancy in Japanese weedy rice, we evaluated the germination response of weedy rice strains described in Table S1. The experiments were repeated four times from 2017 to 2020. Seeds of all strains germinated more when incubated at 30°C than at 15°C. All strains collected in 2017 and 2018 reached almost 100% seed germination at 30°C at 10 weeks after harvest, but most of them failed to reach full germination in 2019 and 2020. Therefore, we conducted another germination assay at 20 weeks after harvest to compare seed dormancy among weedy rice types in 2019 and 2020. We analysed the effects of harvest year, genotype, duration of after-ripening and weedy type on seed germination for each incubation temperature. Significant interactions of After-ripening x Type x Temp x Year and After-ripening x Type x Year at both incubation temperatures were detected by analysis of deviance (Table S4); therefore, we assessed the germination of seeds for each incubation temperature and year.

SH_TRJ showed higher germination rates than BH and SH_TEJ at all time points, incubation temperatures and years (Figure 1, Table S5). For example, for freshly harvested seeds (one week after harvest) incubated at 30°C, the final germination rates of SH_TRJ seeds were over 80% in 2017 and 2018, whereas those of BH and SH_TEJ seeds were lower than 60%. The SH_TRJ seeds in 2019 and 2020 had lower final germination rates than those in the two previous years, but they still showed higher values than the other weedy rice types. SH_TRJ released seed dormancy faster than the other two weedy rice types across all years and incubation temperatures that we tested. The time required to reach 50% final germination rates (T50s) of SH_TRJ seeds were under 10 weeks (or close) in all years at the less permissive temperature (15°C) (Table 1). The T50 of BH weedy rice seeds was 7.67 weeks (7.18 - 8.17) after harvest in 2017 when the seeds were incubated at 15°C but at least 10 weeks in the other years. SH_TEJ weedy rice seeds did not reach T50 before 10 weeks after harvest in all years at 15°C. When incubated in an inductive environment (30°C), SH_TRJ seeds also reached T50 faster than those of other weedy rice types; the 95% CI did not overlap with those of other types except in 2018, showing a significant effect on germination (Table 1).

**FIGURE 1.**
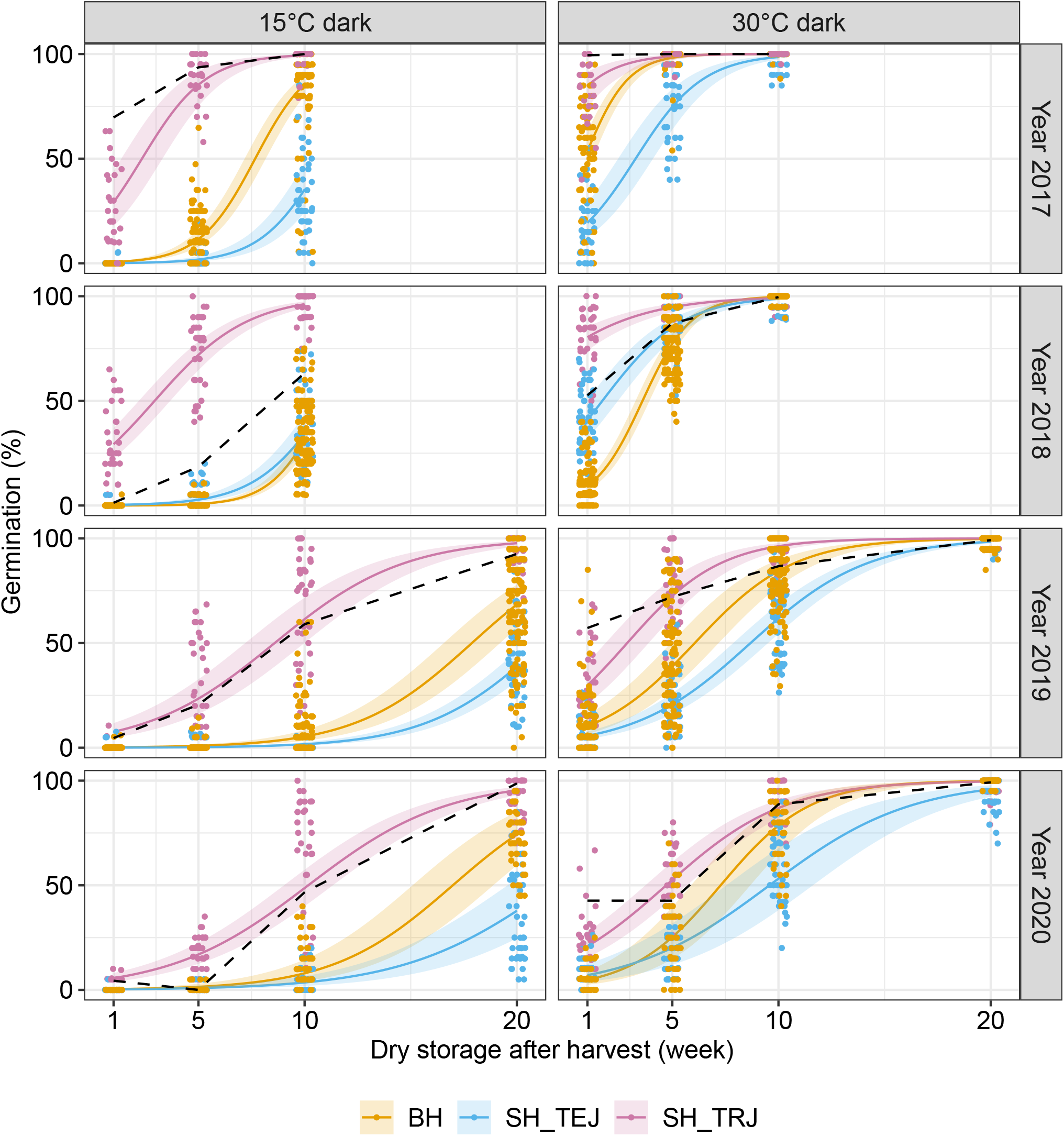
Natural variation for seed dormancy in different weedy rice groups. Degrees of seed dormancy were evaluated by germination rates after different periods of dry storage (x-axis). Germination rates were estimated by a binomial generalized linear mixed model, and strains were used as a random effect. Solid lines represent the maximum likelihood estimates for each strain, and shaded areas represent the 95% confidence intervals for each estimate. Each point indicates germination rates for each strain and replicates at each after-harvest time point. Dashed lines represent the mean germination rates of the cultivated rice variety Koshihikari.

**TABLE 1.** Times to reach 50% final germination rates (T50s) during after-ripening.

|  | Year 2017 | Year 2018 | Year 2019 | Year 2020 |
| --- | --- | --- | --- | --- |
| **15°C dark** |  |  |  |  |
| BH | 7.67 (7.18 – 8.17) | > 10 weeks | 17.88 (16.56 – 19.21) | 16.98 (15.09 – 18.89) |
| SH_TEJ | > 10 weeks | > 10 weeks | > 20 weeks | > 20 weeks |
| SH_TRJ | 2.34 (1.37 – 3.30) | 2.93 (2.19 – 3.67) | 8.59 (7.20 – 9.99) | 10.17 (9.03 – 11.33) |
| **30°C dark** |  |  |  |  |
| BH | < 1 week | 3.60 (3.27 – 3.92) | 6.06 (4.93 – 7.18) | 7.28 (5.89 – 8.67) |
| SH_TEJ | 3.27 (2.45 – 4.09) | 1.70 (< 1 week – 2.44) | 8.69 (7.87 – 9.51) | 9.58 (7.55 – 11.62) |
| SH_TRJ | < 1 week | < 1 week | 2.88 (1.86 – 3.88) | 4.71 (3.74 – 5.69) |

*Note.* Table shows T50 values in weeks and the 95% confidence intervals for seeds of each weedy rice strain incubated either at 15°C or 30°C. The T50 values and the confidence intervals are estimated by a binomial generalized liner mixed model.

SH_TEJ showed deeper seed dormancy than BH (Figure 1). For after-ripened seeds (10 weeks after harvest) incubated at 15°C, the final germination rates of BH seeds were 85.50% (79.98 - 89.69) in 2017, while SH_TEJ seeds germinated lower than 40% that year (Table S5). In 2018, germination of both, BH and SH_TEJ seeds, was similar and below 40%. In 2019 and 2020, BH and SH_TEJ seeds after-ripened for 20 weeks germinated similarly to 10 weeks after-ripened seeds in the previous years when incubated at 15°C: the final germination rates of BH seeds were around 70%; while for SH_TEJ seeds were lower than 40% (Table S5). The T50s of SH_TEJ were consistently higher than those of BH when seeds were incubated at both incubation temperature, except for germination at 30°C in 2018 (Table 1). In 2018, the T50 at 30°C of SH_TEJ was 2 weeks earlier than that of BH: 1.70 week (< 1 week - 2.44) and 3.60 weeks (3.27 - 3.92), respectively.

In summary, the germination assays revealed that SH_TRJ, TRJ-derived weedy rice strains has shallow seed dormancy than BH and SH_TEJ, TEJ derived weedy rice strains. In addition, SH_TEJ, which is derived from the hybridization between weedy and cultivated rice, has deep seed dormancy than the parental weed, BH weedy rice. The differences in seed dormancy levels between BH and SH_TEJ are consistent with our previous study in which we assessed this trait using seeds collected from farmer fields in 2016. In this study, we used seeds collected from an experimental field to remove the effects of the maternal environment caused by provenance. Additionally, we repeated the experiments across a 4-year timeline using seeds collected from weedy rice plants grown in the experimental field from 2017 to 2020. Our results also showed differences in seed dormancy among years: the seeds collected in 2019 and 2020 showed deeper dormancy than those collected in 2017 and 2018 for all weedy rice types suggesting an environmental effect (Figure S1; see Section 3.3 for the description of the results).

### 3.2 Population structure of genomic regions associated with seed dormancy

To assess the population structure of genomic regions associated with seed dormancy among the Japanese weedy rice strains, we performed an admixture analysis based on NGSadmix using a subset of seed dormancy-related regions. The genetic subgroups for the genomic regions at *K* = 5, the best model in the whole genome sequencing analyses (Imazumi et al. 2021), corresponded to the following groups of accessions (Figure 2a): (1) BH and SH_TEJ weedy rice; (2) modern_TEJ and landrace_TEJ; (3) SH_TRJ weedy rice and landrace_TRJ; (4) Chinese weedy rice; and (5) landrace_IND and US weedy rice. The admixture analysis results of SH_TRJ weedy rice strains were consistent with those for the whole genomes. In contrast, admixture analysis results of SH_TEJ weedy rice strains were different between the regions related to seed dormancy and the whole genome sequences. Here, SH_TEJ weedy rice strains belonged to the same group as BH_weedy rice in terms of seed dormancy, but they showed admixed ancestry between BH weedy rice and modern/landrace_TEJ rice from Japan in the whole genome. Evaluation of the likelihood of *K* (*L(K)*, Pritchard et al., 2016) suggested that the likelihood reached a plateau at *K* = 4 or 5, but the peak of Δ*K* (Evanno et al., 2005) was detected at *K* = 2 (Figure S2b and c). Thus, the results from *K* = 2 to 4 are also presented for comparison. Cultivated rice has been classified into two major groups: *japonica* (TEJ and TRJ) and *indica* (IND and AUS). At *K* = 2 and 3, the genetic subgroups correspond to *japonica* and *indica* and to TEJ, TRJ and *indica*, respectively (Figure S2a). At *K* = 4, modern_TEJ and some landrace_TEJ strains are distinct from BH weedy rice, but SH_TEJ weedy rice strains belong to the same group as BH_weedy rice, which is similar to the result at *K* = 5.

**FIGURE 2.**
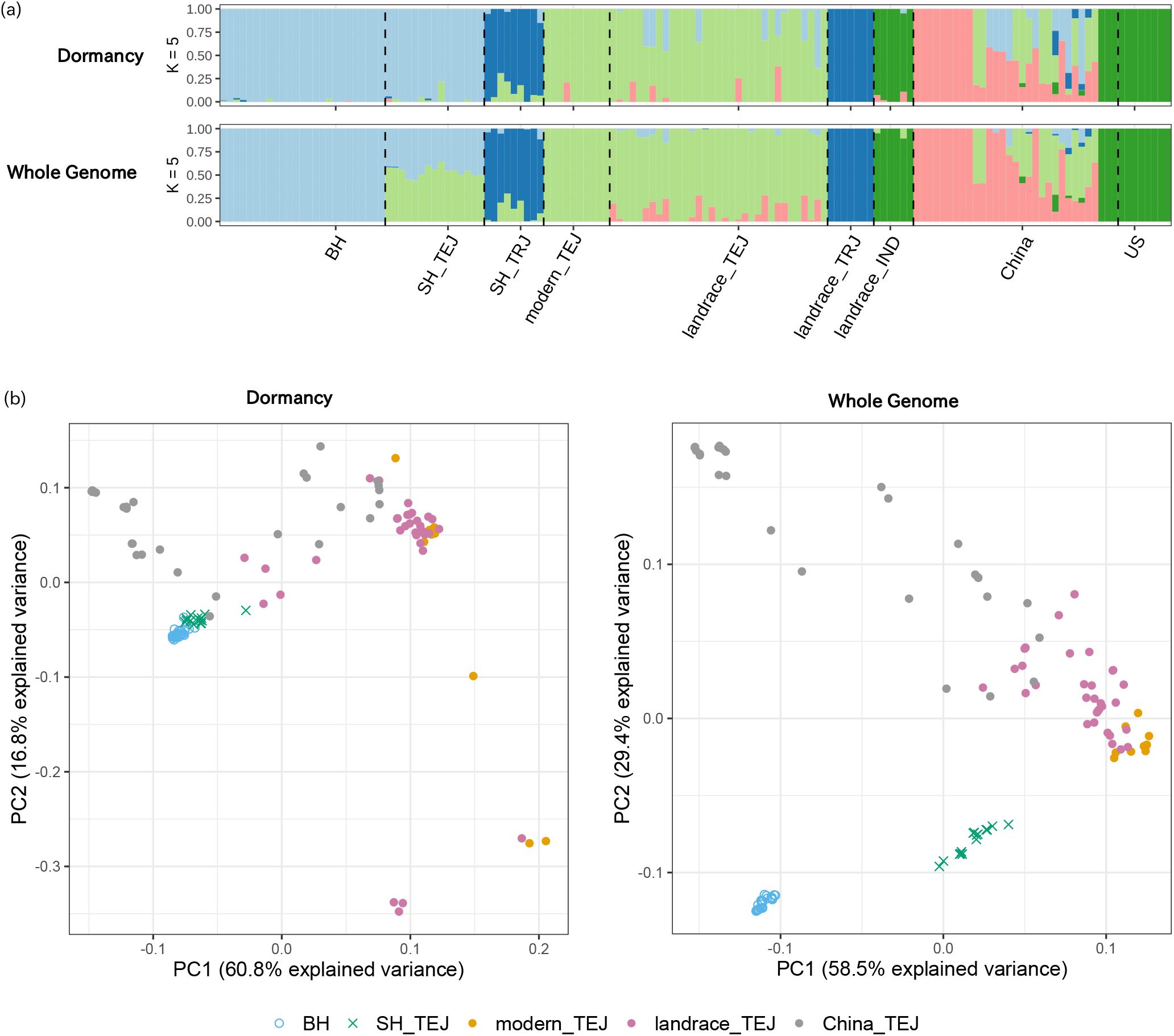
Genomic classification of Japanese weedy rice strains using genomic regions associated with seed dormancy. (a) Population structure of weedy and cultivated rice strains using genomic regions associated with seed dormancy (upper) and whole genome sequence (lower). Ancestry proportions for individuals with *K* = 5 are presented in the plots. (b) PCA plot of 111 TEJ rice strains, including BH, SH_TEJ, modern_TEJ, landrace_TEJ and China_TEJ; the first and second eigenvectors were obtained using genotype likelihoods estimated by ANGSD using the genomic regions associated with seed dormancy (left) and the aligned and mapped reads of whole genome sequencing data (right).

PCA using the variants in the regions related to seed dormancy across weedy and cultivated TEJ rice also supported the close relationship between BH and SH_TEJ weedy rice strains (Figure 2b). In contrast, PCA separated all of the modern TEJ cultivated rice varieties, most of the landrace TEJ cultivated rice varieties, and Chinese weedy rice strains from BH and SH_TEJ weeds, which is consistent with the PCA results obtained using the variants from the whole genome sequences. The first principal component explained 60.8% of the total variation and separated the Japanese weedy rice strains and cultivated rice varieties, and the second principal component explained 16.8% of the total variation, separating the Japanese and Chinese weedy rice strains. Interestingly, some of the landrace varieties and the Chinese weedy rice strains were plotted close to the Japanese weedy rice strains, consistent with the admixture analysis for some landrace_TEJ varieties showing admixed ancestry between BH weeds and modern/landrace_TEJ cultivated rice (Figure 2a and b). PCA plot of all strains used in this study separated SH_TRJ from BH and SH_TEJ weedy rice strains for both of the seed dormancy and the whole genome sequence (Figure S3), which was also demonstrated with the results of admixture analysis.

The admixture analysis and PCA results showed that SH_TEJ weedy rice strains shared most of the genomic regions associated with seed dormancy with BH weedy rice strains, suggesting that SH_TEJ weedy rice maintains the adaptive allelic combinations for seed dormancy derived from BH weedy rice. We summarised the variants detected in BH weedy rice for the 131 genes associated with seed dormancy (Table S3) and compared the variants with the other TEJ weedy and cultivated rice strains in Japan: SH_TEJ; modern_TEJ; and landrace_TEJ (Table S6). Variants with the Nipponbare reference sequence, which are fixed or mostly fixed in BH weedy rice, were detected for 36 genes, and the variants were located on all chromosomes, except chromosome 9. SH_TEJ weedy rice strains shared 69.4% of the fixed alleles with BH strains (25 from the 36 alleles). Among the 25 alleles, fixed or mostly fixed alleles within SH_TEJ weedy rice strains were observed for 22 genes. SH_TEJ weedy rice has two subgroups (SH1_TEJ and SH2_TEJ, Imaizumi et al., 2021): SH1_TEJ had the same alleles as BH of the genes *OsBT1-1* (Os02g0202400), *OsPLL3* (Os02g0214400) and *OsLHT1* (Os08g0127100), whereas these were different for SH2_TEJ (Table S6). For the gene *SAP16* (Os07g0569700), all SH_TEJ weedy rice strains, except for JP_1162, had the same allele as BH. In contrast, no loci had the same fixed allele as BH in modern and landrace varieties, except for *OCP* (Os04g0650000) in modern varieties and *OsFY* (Os01g0951000) in landrace varieties.

### 3.3 Differences in seed dormancy among years and the effects of maternal temperature

The germination rates were generally lower in the 2019 and 2020 experiments compared with the 2017 and 2018 experiments (Table 1; Figure S1). The daily average temperature after heading (from August to September) was higher in 2019 and 2020 than in 2017 and 2018 (Figure S3), indicating that maternal temperature may affect seed dormancy in the progeny. To evaluate this possibility, we calculated the cumulative daily mean temperature for 40 days after heading. The maternal effects were evaluated using the germination rates of 5-week after-ripened seeds incubated at 30°C and 10-week after-ripened seeds incubated at 15°C because the combinations of those time points and germination conditions accurately explained the differences in seed dormancy among years (Figure S1). Statistical assessment of the maternal effects revealed that higher temperatures after heading significantly decreased the germination rates in all weedy rice strains (Figure 3), suggesting that higher temperatures increased progeny dormancy. For example, for every 20°C of temperature that accumulated daily in average, the odds of germination increased by 0.65 (0.63-0.66) for BH seeds that were after-ripened for 10 weeks and incubated at 15°C (Table 2).

**FIGURE 3.**
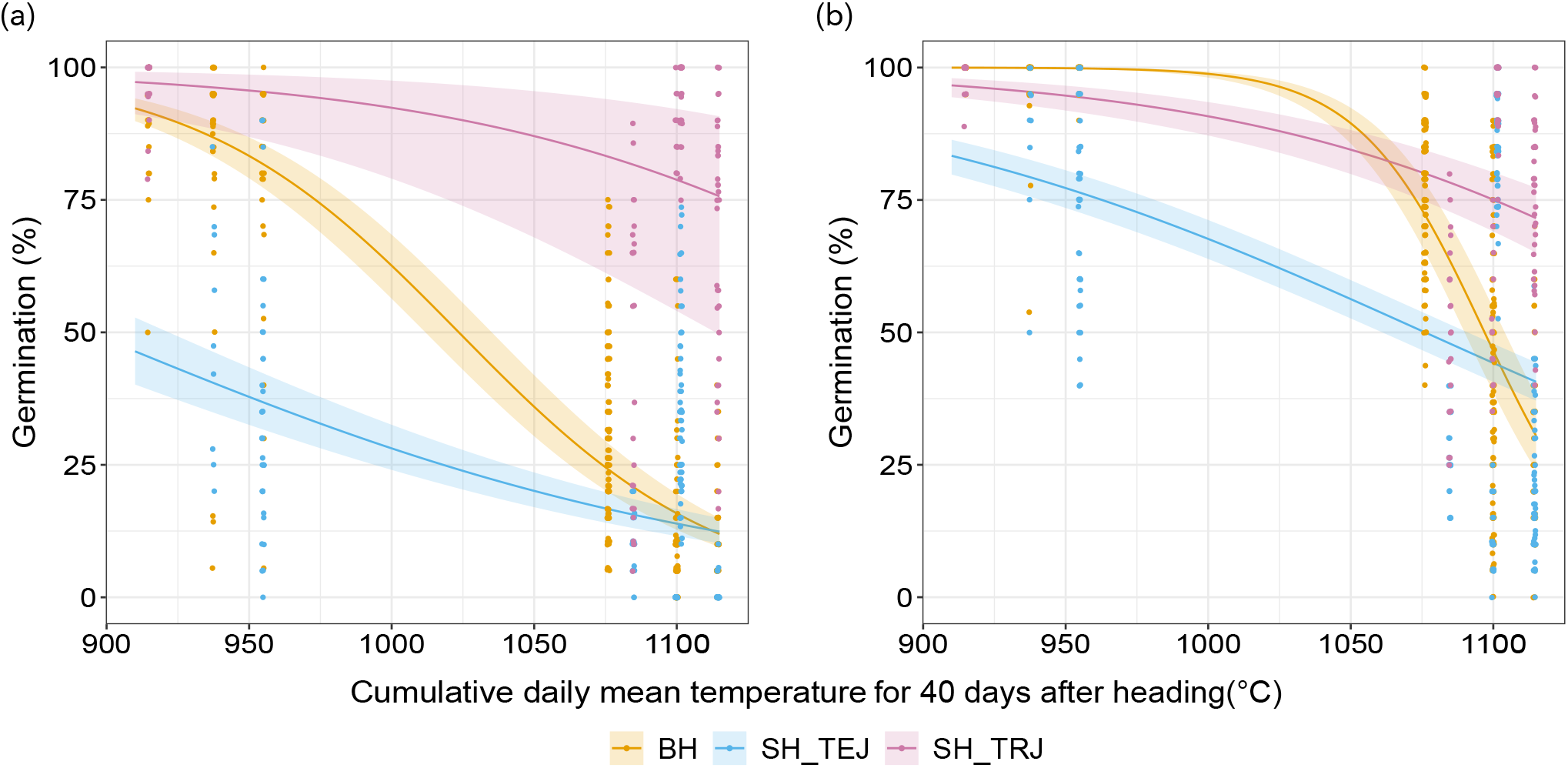
Maternal environment effects on progeny seed dormancy in weedy rice. Effect of cumulative daily mean temperature after heading on progeny seed dormancy evaluated by germination rates of (a) 10-week-old seeds incubated at 15°C and (b) 5-week-old seeds incubated at 30°C. Germination rates were estimated by a binomial generalized linear mixed model, and strains were used as a random effect. Shaded areas represent the 95% confidence intervals for each curve, and each point indicates germination rates for each strain and replicates at each post-harvest time point.

**TABLE 2.** Odds ratios for the effect of cumulative daily mean temperature after heading on progeny seed dormancy.

|  | 10 weeks AR seeds<sup>1)</sup> |  | 5 weeks AR seeds<sup>2)</sup> |  |
| --- | --- | --- | --- | --- |
|  | odds ratio | 95% CI | odds ratio | 95% CI |
| BH | 0.65 | 0.63 – 0.66 | 0.4 | 0.37 – 0.43 |
| SH_TEJ | 0.84 | 0.82 – 0.86 | 0.82 | 0.81 – 0.84 |
| SH_TRJ | 0.79 | 0.75 – 0.83 | 0.79 | 0.75 – 0.83 |

*Note.* Odds ratios estimated by a binomial generalized liner mixed model are presented.
AR = after-ripened; CI = confidence interval;
<sup>1)</sup> results for germination rates under 15°C in the dark for 10-week-old AR seeds;
<sup>2)</sup> results for germination rates under 30°C in the dark for 5-week-old AR seeds.

For BH and SH_TEJ weedy rice, the maternal temperature effects were also confirmed using greenhouse experiments. Progeny dormancy was higher for both BH and SH_TEJ weedy rice if plants were grown under a diurnal temperature cycle of 34°C/28°C after heading than when grown at 28°C/22°C (Figure S4). In general, the germination response of freshly harvested seeds (1 week after harvest) was similar between seeds matured under both treatments. However, in some instances, seeds that matured under the high-temperature treatment showed higher germination rates immediately after harvest, for example, BH seeds incubated at 30°C in 2016 and SH seeds incubated at 15°C and 30°C in 2018. However, seeds after-ripened for 10 weeks obtained from mother plants grown under the high-temperature treatment germinated less than those from control mother plants (Figure S4). Our results suggest that maternal temperature affects dormancy breakage and not primary dormancy of progeny seeds under our conditions.

## 4 DISCUSSION

In this study, we found variation in seed dormancy among weedy rice strains, population structure in genomic regions associated with seed dormancy, and maternal temperature effects. We used three weedy rice types to evaluate the variation in seed dormancy: TEJ-derived blackhull and strawhull (BH and SH_TEJ, respectively) and TRJ-derived strawhull (SH_TRJ) weedy rice. We found that SH_TRJ expresses shallow seed dormancy, BH has an intermediate level of seed dormancy, and SH_TEJ shows deep seed dormancy. In general, BH ecotypes are more dormant than SH ecotypes (Diarra et al., 1985; Tseng et al., 2013), and the black pigmentation of the hull also plays an important role in imposing seed dormancy (Diarra et al., 1985; Gu et al., 2005). However, previous studies compared the seed dormancy of BH and SH weedy rice derived from different cultivated rice groups, such as comparing BH weedy rice derived from *aus* to SH weedy rice derived from *indica* (Tseng et al., 2013, 2018). When we compared the BH and SH ecotypes derived from TEJ cultivated rice, the SH weedy rice had deeper seed dormancy than the BH weedy rice (Figure 1), indicating that phylogenetic constraints for the degree of seed dormancy should be considered in addition to the effects of BH pigmentation.

SH_TEJ weedy rice showed the deepest seed dormancy among the weedy rice types evaluated in this study although the weeds were derived from hybridization with cultivated rice (Imaizumi et al., 2021), which must be a non-adaptive parent for weedy-like traits, including seed dormancy. Seed dormancy in cereal crops should be moderate because high seed dormancy impairs the establishment of crops, and a lack of seed dormancy causes preharvest sprouting, significantly reducing the yield and quality (Rodríguez et al., 2015; Tuan et al., 2018). Seed dormancy is associated with plant height in rice: semidwarf genotypes have deeper seed dormancy than tall genotypes based on variation in the genes *semidwarf 1* (*sd1*) and *ent-kaurene oxidase* (*KO*), which are associated with gibberellin (GA) biosynthesis and that regulate seed dormancy and plant height in rice (Ye et al., 2013; Wang et al., 2020; Itoh et al., 2004; Wu et al., 2014; Zhang et al., 2020). Our previous study detected genetic contributions of introgressed alleles for seed dormancy at the *KO1* and *KO2* loci from cultivated rice (Imaizumi et al., 2021). In addition, this study revealed that the hybrid-derived weedy rice strains maintained most of the seed dormancy alleles that were observed in the parental weedy rice (Figure 2 and Table S6). Therefore, the deep seed dormancy in SH_TEJ weeds is most likely derived from both BH weedy rice (the adaptive allelic combinations of seed dormancy-related genes) and cultivated rice (the adaptive introgression of deep seed dormancy traits).

Hybridization is a common evolutionary process; however, many unanswered questions still exist regarding its genetic and evolutionary mechanisms (Moran et al., 2021). One question is whether genome stabilization after hybridization is typically driven by genetic drift or selection. The present study shows that the stabilization of genomic regions associated with seed dormancy occurred by the removal of alleles derived from parental cultivated rice in hybrid-derived weedy rice (Figure 2 and Table S6). Whole genome sequence analyses detected signatures of positive selection on some regions regulating seed dormancy in hybrid-derived weedy rice and parental weedy rice (Imaizumi et al., 2021). Therefore, genome stabilization for the regions associated with seed dormancy would be driven by positive selection in hybrid-derived weedy rice, and most of the alleles associated with this trait are fixed or nearly fixed for parental weedy rice. Most of the genomic regions derived from the minor parent were eliminated after hybridization (Schumer et al., 2016; Matute et al., 2020; Chaturvedi et al., 2020), suggesting positive selection against the minor parent. In contrast, the hybrid-derived weedy rice used in this study was not highly biased toward the parental weedy rice but showed the genome composition combined in a mosaic in the whole genome (Figure 2a, Imaizumi et al., 2021). This study highlights genome stabilization by positive selection for seed dormancy, a major life-history trait crucial for local adaptation (Donohue et al., 2010). This is supported by the results showing that most of the alleles associated with seed dormancy are fixed or nearly fixed regardless of the mosaic composition in the whole genome.

The yearly variation in seed dormancy suggested an environmental cue that influenced the expression of variation for this trait in the genotypes analysed in this study, revealing that maternal environments influences variation in seed dormancy among weedy rice strains. Maternal temperature during seed maturation is a strong signal across species as an environmental factor for generating variation in seed dormancy (Peters, 1982; Reddy et al., 1985; Burghardt et al., 2016; Penfield & MacGregor, 2017). We observed effects of maternal temperature after heading on progeny seed dormancy: seeds that matured under higher temperatures had deeper seed dormancy. Under certain conditions, maternal effects were stronger than genetic effects. For example, the estimated germination rate of BH seeds matured under warmer conditions (27.5°C daily mean temperature) was lower than that of SH seeds matured under cooler environments (23.75°C daily mean temperature) (Figure 3). The odds of germination for the increasing temperature were lower for BH seeds than for SH_TEJ seeds (Table 2), suggesting BH weedy rice was more sensitive to the maternal effects. High temperature after heading reduces expression of ABA catabolism genes in rice seeds (Suriyasak et al., 2020). A key gene in ABA catabolism in rice seeds is *OsABA8ox3* (Zhu et al., 2009), and BH weedy rice has a different allele in *OsABA8ox3* compared with SH weedy rice (Table S6). This allelic variation may influence the difference in sensitivity to the maternal effects between the two weedy rice types.

This study also revealed that the genetic structure of genomic regions associated with seed dormancy is constrained by phylogeny as genome structure differed between TEJ-derived and TRJ-derived weedy rice strains. Natural variation for seed dormancy in *A. thaliana* is mainly controlled by different additive genetic and molecular pathways rather than epistatic interactions, indicating the involvement of several independent pathways (Bentsink et al., 2010). Genomic analyses in weedy rice indicate that *GD1*, a gene associated with seed germination, is a shared, strong candidate for adaptive traits in TEJ-derived weedy rice from different countries, such as Japan, China, South Korea and Italy (Qiu et al., 2020). However, this gene is missing in TRJ-derived weedy rice in Japan (Imaizumi et al., 2021). We also observed that different weedy rice strains originating from independent feralization events show differences in genomic structure in regions associated with seed dormancy: TEJ-derived weedy rice strains (BH and SH_TEJ weeds) were clustered in a different group from SH_TRJ.

Most weedy rice strains regardless of their origin (TEJ TRJ, IND or AUD) share the gene *Rc* as determinant for seed dormancy. The functional allele *Rc* from weedy red rice was observed to play a major role in maintaining seed morphological integrity and viability in the field; however, a single gene, including *Rc*, did not appear to provide a sufficient degree of dormancy and longevity for seeds to survive in the soil from the winter to spring seasons (Pipatpongpinyo et al., 2020). At least 30 QTLs have been associated with seed dormancy, each of the QTLs alone explain a low proportion of the phenotypic variation in this trait, indicating a high complexity in its regulation (Wang et al., 2018). The different genetic backgrounds of genomic regions associated with seed dormancy among Japanese, Chinese and US weedy rice strains suggest the possibility of a wider range of variation in seed dormancy worldwide.

## 5 CONCLUSION

Our phenotypic and genomic analyses of seed dormancy indicate a robust variation mechanism within species. Different weedy rice strains with independent origins show variation in genomic structure for seed dormancy, which suggests the possibility of local variation in this trait in weedy rice worldwide. Local adaptation typically requires fixation of alleles at multiple loci that contribute to increased fitness in a certain environment. Our analyses suggest that the history of hybridization between weedy rice (having adaptive allelic combinations for seed dormancy) and cultivated rice (having non-adaptive combinations) has generated weedy rice strains that express deep seed dormancy caused by genome stabilization through the removal of alleles derived from cultivated rice, in addition to the adaptive introgression of deep seed dormancy alleles derived from cultivated rice.

## ACKNOWLEDGEMENTS

The authors thank the staff members of Weed Management Group, Institute for Plant Protection, NARO and those of the Department of Technical Support of NARO for their technical assistance. This research was supported by the Advanced Analysis Center Research Supporting Program of National Agriculture and Food Research Organization (NARO) and the Advanced Genomics Breeding Section of Institute of Crop Science, NARO (NICS). We also thank the Advanced Analysis Center of NARO (NAAC) and AFFRIT of Ministry of Agriculture, Forestry and Fisheries (MAFF), Japan for use of the high performance cluster computing system, respectively. The authors thank Kentaro Ohigashi (NARO) for helpful comments on statistical analysis. This work was supported by grants from the Project of the Bio-oriented Technology Research Advancement Institution, NARO (the special scheme project on advanced research and development for next-generation technology), and from the commissioned project “Development of labor-saving management of serious weeds to expand cultivation of direct-seeded rice”, MAFF, Japan.

## AUTHOR CONTRIBUTIONS

T.I. designed and managed the research. T.I. and Y.K. performed the bioinformatic analyses. T.I.and G.A. carried out the experiments. T.I. wrote the original draft of the manuscript. All authors revised, read and approved the final manuscript.

## BENEFIT-SHARING

Benefits Generated: Benefits from this research accrue from the sharing of our data and results on public databases as described above.

## DATA AVAILABILITY STATEMENT

Short-read sequence data of 144 samples used in by this study can be downloaded from DDBJ/EMBL/GenBank with the BioSample accesion numbers described in Table S2.

