## Supplementary Figures for "Hybrid-derived weedy rice maintains adaptive combinations of alleles associated with seed dormancy"

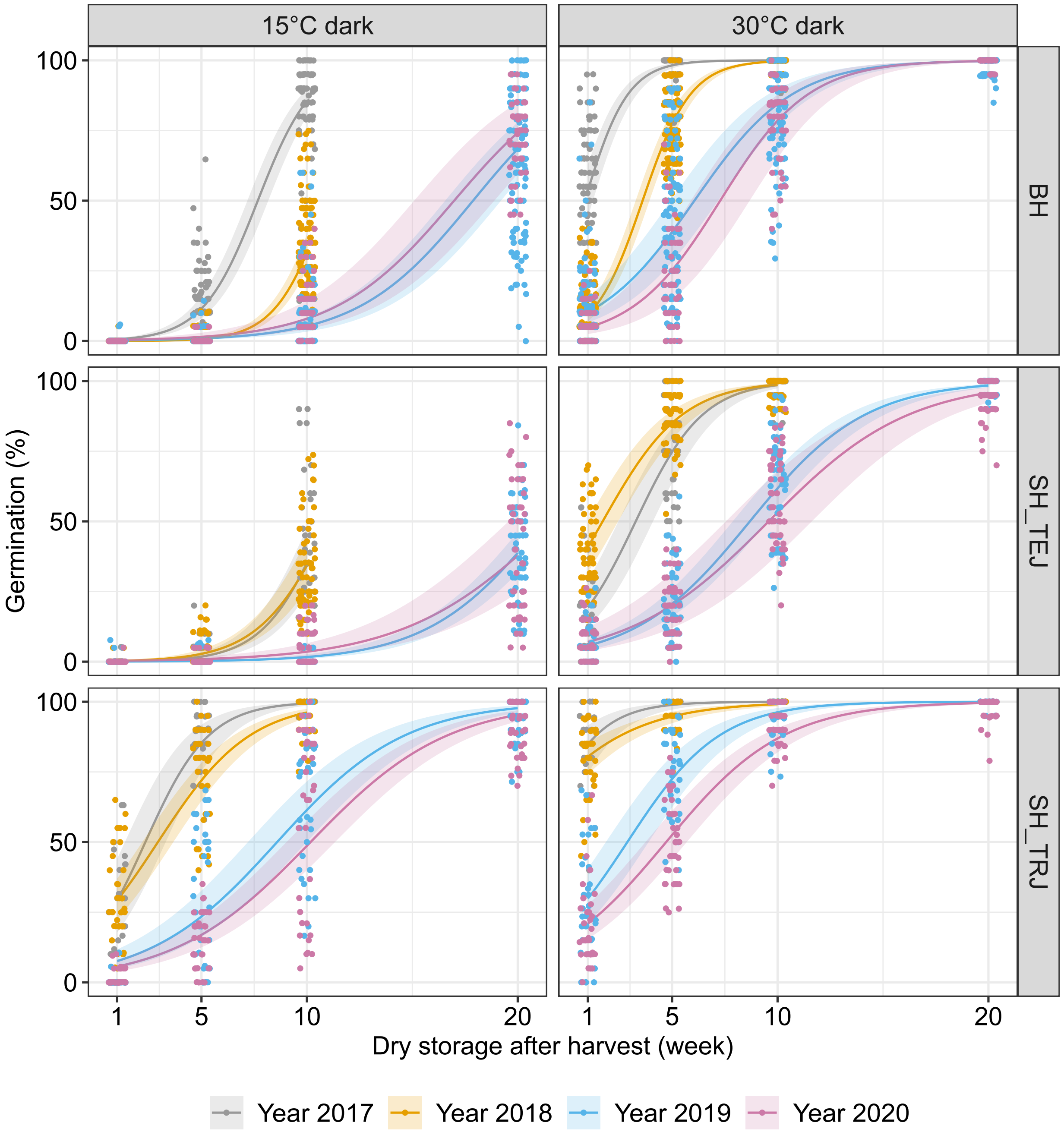


**FIGURE S1** Variation in seed dormancy among different years for each weedy rice type. Graphs show the germination rates by weedy rice type (rows) and incubation conditions (columns). The germination rates from Figure 1 were used to compare the variation in seed dormancy among the years. Solid lines represent the maximum likelihood estimates for each year, and shaded areas represent the 95% confidence intervals for each estimate. Each point indicates germination rates for replicates in a given year and after-harvest time point.

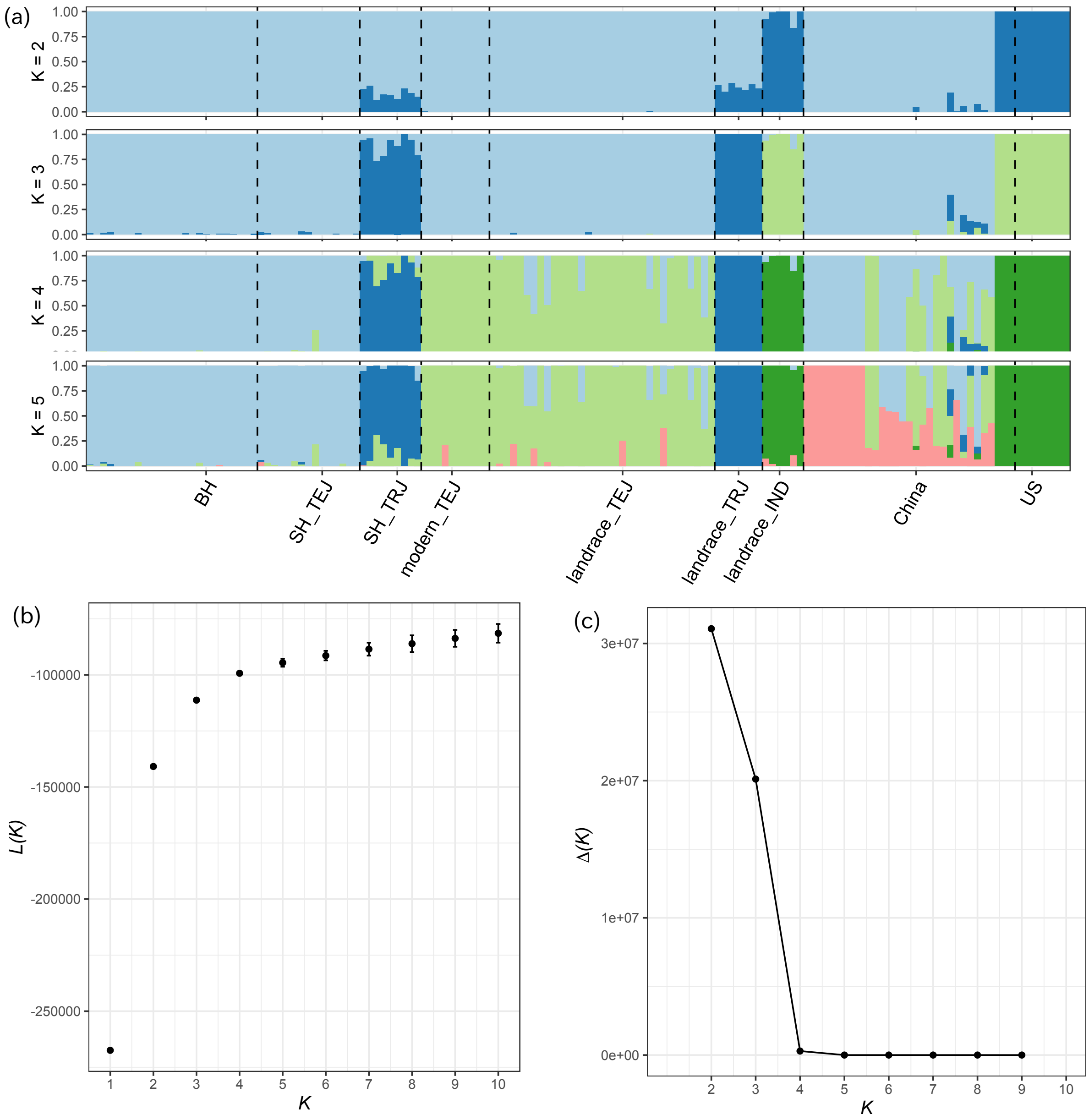


**FIGURE S2** Optimal number of $K$ values evaluated by $L(K)$ and $\Delta K$ values. (a) Population structure of weedy and cultivated rice strains using genomic regions associated with seed dormancy. Ancestry proportions for individuals with K = 2 to 5 are presented in the plots. (b) Mean $L(K)$) (±SD) over 10 runs for each $K$ value up to $K = 10$. (c) $\Delta K$ values for each $K$ value up to $K = 9$.

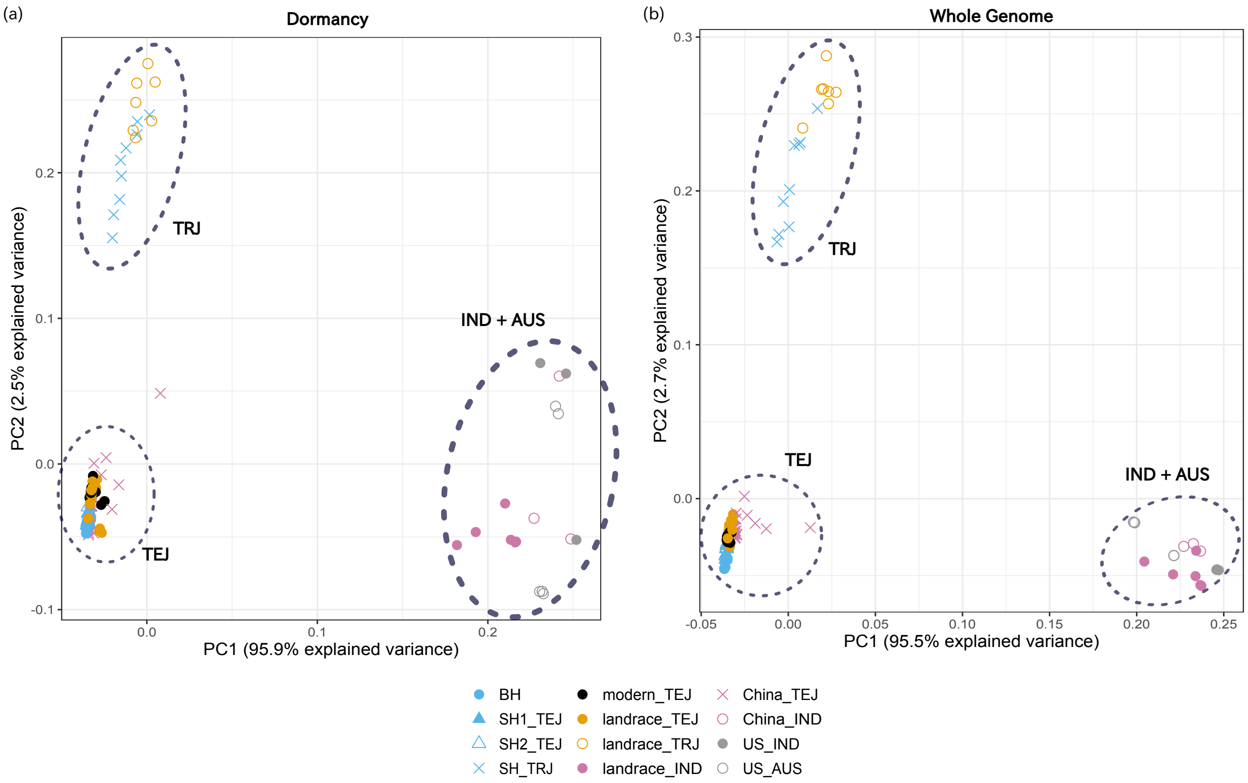


**FIGURE S3** PCA plot of all strains used in this study. The first and second eigenvectors were obtained using genotype likelihoods estimated by ANGSD using the genomic regions previously reported as associated with seed dormancy (a) and the aligned and mapped reads of whole-genome sequencing data (b).

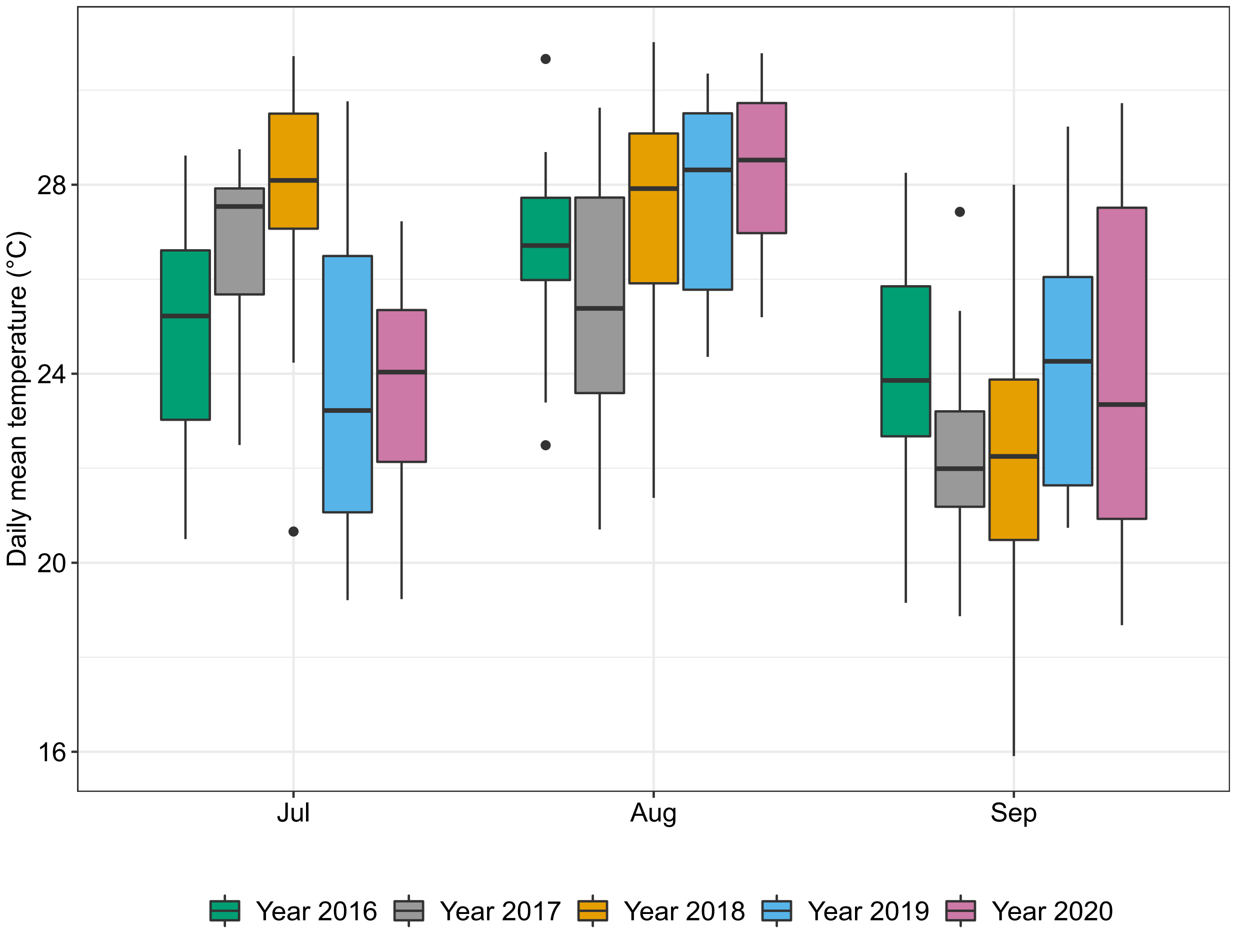


**FIGURE S4** Box plot of the daily mean temperature for July, August and September at the Tsukuba-Kannondai test field, recorded by the Weather Data Acquisition System of the Institute for Agro-Environmental Sciences, NARO, between 2016 and 2020.

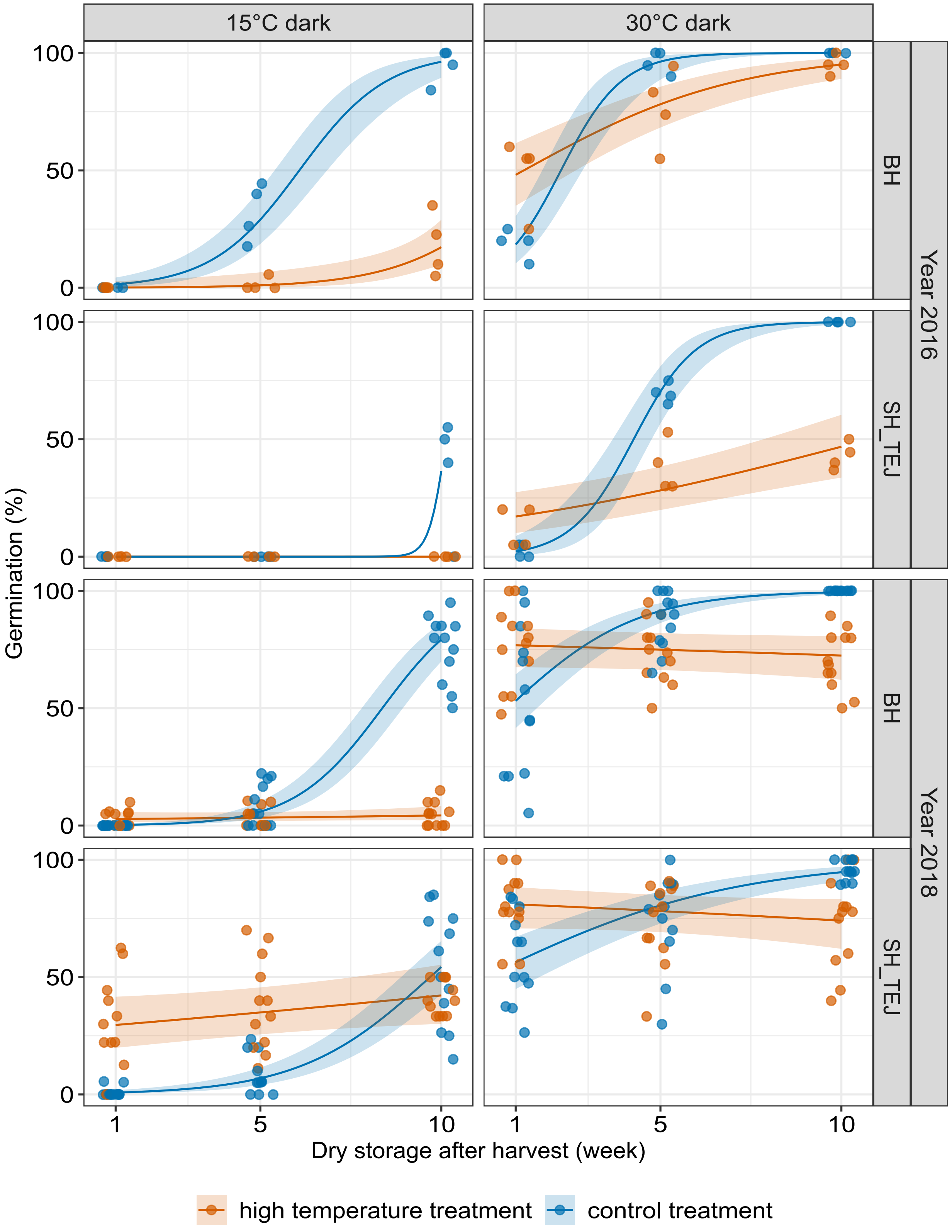


**FIGURE S5** Effect of maternal temperature after heading on progeny seed dormancy. Germination rates of progeny seeds incubated at 15 or 30°C during an after-ripening time course for two experiments run in 2016 and 2018. Orange lines represent germination rates of seeds matured under the high-temperature treatment (34°C/28°C), blue lines represent those of seeds matured under the control treatment (28°C/22°C), and shaded areas represent the 95% confidence intervals for each curve. Each point indicates germination rates for each replicate at every time point.
